# Quantum Machine Learning for Missense Variant Pathogenicity Prediction in the KCNQ Gene Family

**DOI:** 10.64898/2026.11.16.767028

**Authors:** Markel G. Ibarluzea, Sara Capponi, Aitor Bergara, Aritz Leonardo

## Abstract

Missense variants in the KCNQ family of voltage-gated potassium channel genes are associated with a broad spectrum of neurological and cardiac disorders for which computational tools capable of differentiating benign and pathogenic variants are essential. Here, we present, to our knowledge, the first application of quantum machine learning (QML) to missense pathogenicity prediction. We employ a hybrid quantum framework combining data re-uploading quantum neural networks with quantum support vector machines and applied it to pathogenicity classification across the full KCNQ gene family. Systematic exploration of circuit architectures revealed that optimal hyperparameter choices enable the proposed quantum model to achieve predictive performance superior to or comparable to state-of-the-art classical machine learning approaches, reaching AUC-ROC values of approximately 0.94 across the full dataset. Although no clear quantum advantage is observed at present, these results demonstrate that quantum feature representations can effectively capture biologically relevant patterns in variant data and establish QML as a viable framework for future applications in computational genomics and precision medicine.

## Introduction

Missense variants in voltage-gated potassium channel genes represent a broad and clinically heterogeneous category of genetic pathology, encompassing conditions that range from benign, self-resolving neonatal seizures to severe, intractable developmental and epileptic encephalopathies. ^1–4,4^ The KCNQ gene family, which encodes the KV7 subfamily of voltage-gated potassium channels, is a great example of this clinical diversity. ^5–7^ Accurate early classification of variant pathogenicity and anticipated phenotypic severity is essential to guide individualized therapeutic decisions and genetic counseling across this entire gene family. The rapid expansion of genomic sequencing technologies has greatly increased variant discovery, ^8^ but the task of distinguishing genuinely pathogenic variants from tolerated polymorphisms, and of further stratifying pathogenic variants by expected clinical severity, remains an unresolved challenge. ^9^

Computational tools for variant effect prediction have emerged as a critical resource for interpreting missense mutations at scale, with both traditional machine learning based techniques, ^10–13^ as well as models based on deep learning foundation models, ^14–16^ showing promising performance. Despite their improving performance, these tools still underperform for specific gene families, and their accuracy has been shown to be highly gene-dependent.^17^ For the KCNQ family in particular, their performance varies by member, with the best performing models approaching clinically acceptable performance for KCNQ1 and 2, while struggling with the other 3 family members. Our prior work with on a predictive model for KCNQ2 showed that gene-specific models can improve accuracy over generic tools; ^18^ the present study extends this framework to the full KCNQ family, recognizing that the shared topology and functional conservation of KV7 channels create a natural basis for a unified, family-level model.^19^

In this work, we analyze the potential of quantum machine learning, a nascent field whose potential applications in life sciences are of growing interest, ^20–22^ to overcome some of the limitations of current variant effect prediction methods. A key conceptual motivation for QML in biological applications is that genomic and proteomic data are inherently high-dimensional and structurally complex, characteristics that could potentially favor quantum representations over classical ones. ^23^ Recent studies demonstrate that quantum methods can match or exceed classical benchmarks in tasks like analyzing electronic health records, genomic sequences, and immunotherapy data. ^24,25^ However, the field remains nascent, with further hardware and algorithmic advances needed to unlock QML’s full potential in clinical bioinformatics.

Building on these advances, this study presents what is, to our knowledge, the first application of quantum machine learning to pathogenicity classification of missense variants. We develop a QML model that integrates quantum kernel and data re-uploading approaches, along with the gene-family-specific feature engineering strategy established in our prior classical work, and test a broad range of model settings and hyper parameters to assess the configurations under which QML models show optimal performance. The methodology presented here, which combines gene-family-level biological priors with quantum feature representations, offers a transferable framework for variant interpretation in other ion channel gene families and, more generally, for the application of QML to rare disease genomics.

## Methods

### Dataset and Feature Representations

We constructed a dataset of missense variants from members of the KCNQ ion channel gene family with data obtained from the publicly available databases ClinVar, ^26^ RIKEE and LOVD3. ^27^ Variants annotated as pathogenic or likely pathogenic were combined as the pathogenic class, whereas those annotated as benign or likely benign were used as the benign class. Because ClinVar often contains imbalanced data that is biased towards pathogenic variants, due to physicians naturally having higher likelihood of testing patients with some kind of pathogenicity, we supplemented the benign class using population frequency data from Genome Aggregation Database (gnomAD). ^28^ Although gnomAD does not contain labeled data, certain variants are observed at appreciable frequencies in healthy populations. These variants were assumed to be benign and incorporated as negative examples. This combined strategy increases the availability of confidently labeled benign variants while preserving clinically validated pathogenic annotations.

Each variant was represented using a set of features designed to capture complementary biological information relevant to protein function and disease association. These features fall into three broad categories. First are the features that describe the structural context of the mutated residue, including the predicted secondary structure and the local confidence of the predicted structure as measured by the pLDDT score produced by AlphaFold2. ^29^ Secondly, physicochemical features that characterize the effect of the amino acid substitution on key residue properties such as size, polarity, hydrophobicity, charge, and side-chain volume. Finally, evolutionary features that capture the degree of conservation at the mutated position within the protein sequence, reflecting the evolutionary constraints acting on that site.

### Quantum Machine Learning Model

The quantum machine learning model used in this work is based on the architecture introduced in, ^30^ which combines a quantum neural network (QNN) with a quantum support vector machine (QSVM). The QNN is first trained as a binary classifier and its optimized circuit is subsequently fixed and used as the quantum feature map from which the kernel of the QSVM is computed.

The QNN follows a data re-uploading architecture, ^31^ consisting of repeated layers of data-encoding unitaries, trainable single-qubit unitaries, and entangling operations. For an input triplet **x** = (*x*_1_, *x*_2_, *x*_3_), the data-encoding operation applied to each qubit is

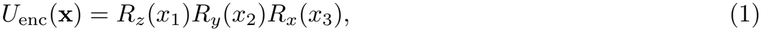

If fewer than three input components are available, the corresponding trailing rotations are omitted. The trainable operation acting on qubit *q* in layer *l* follows the same composition

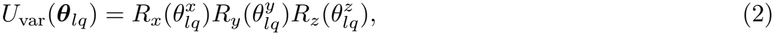

For an *n*-qubit circuit, the unitary associated with layer *l* can therefore be written as

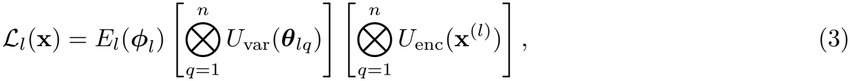

where *E*_*l*_(***ϕ***_*l*_) denotes a sequence of parameterized controlled-*SU* (2) operations between adjacent qubits. Entangling operations are inserted between consecutive data-reuploading layers and are omitted after the final layer. The complete circuit is

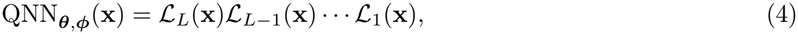

with the layer-dependent input **x**^(*l*)^ determined by the embedding strategy described below.

Only the first qubit is measured during QNN training. The probability of obtaining measurement outcome 1 is interpreted as the predicted probability of the pathogenic class. The circuit parameters are optimized by minimizing a binary cross-entropy loss using the Adam optimizer with an initial learning rate of 0.1. Training is performed in mini-batches of 64 samples. The learning rate is multiplied by 0.95 after each epoch. The initial single-qubit circuit is trained for 30 epochs, whereas each subsequent circuit obtained by adding a new qubit is trained for 10 epochs.

Circuit training proceeds iteratively, with qubits introduced one at a time. After training the single-qubit circuit, its optimized parameters are transferred to the corresponding operations in a two-qubit circuit. The parameters belonging to the newly introduced qubit and its parameterized entangling operations are initialized to zero. Because these zero-valued operations reduce to the identity and only the first qubit is measured, the expanded circuit initially reproduces the input-output mapping and loss value of the previously optimized circuit. Training therefore resumes from the solution obtained at the preceding circuit size while introducing additional variational degrees of freedom. This procedure is repeated until the target number of qubits is reached.

Once QNN training is complete, the optimized parameters (***θ, ϕ***) are fixed and the resulting circuit is repurposed as a quantum feature map. Kernel entries for two input samples **x**_*i*_ and **x**_*j*_ are calculated from the state fidelity

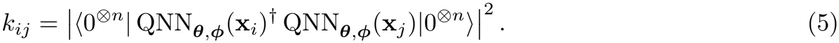

The resulting Gram matrix is then supplied as a precomputed kernel to a support vector machine, which performs the final pathogenicity classification.

### Data Embedding

Before quantum embedding, the input features were standardized and reduced in dimensionality using principal component analysis (PCA), and the top *d* components with highest explained variance were retained as inputs to the QNN, were they were applied directly as rotation angles in the data-encoding unitaries. Both transformations were fitted exclusively on the training data and subsequently applied without refitting to the held-out data.

A single *SU* (2) encoding block accommodates at most three principal components, each parametrizing a rotation in each of the 3 possible axes. The same encoded component values are applied to every qubit. To incorporate more than three components, we evaluated two embedding strategies.

In the stacked strategy, the retained components are divided into consecutive groups of at most three. All groups are encoded sequentially within every circuit layer. Thus, each layer contains multiple encoding blocks and receives the complete retained feature representation.

In the cyclic interleaving strategy, the retained components are likewise partitioned into groups of at most three, but successive groups are assigned to successive circuit layers. If the number of layers exceeds the number of groups, assignment cycles back to the first group. For example, with nine retained components, layers one, two, and three encode components (1, 2, 3), (4, 5, 6), and (7, 8, 9), respectively, after which the sequence is repeated in later layers. This distributes the input representation across the circuit depth rather than applying all components within every layer. The number of retained components is therefore constrained to at most three times the number of circuit layers. The cyclic interleaving strategy was used for the final models.

## Results

We evaluated the quantum-kernel model using feature representations that combined structural, physicochemical, evolutionary, and gene-family information. The principal component representation, circuit architecture, and model-training procedure were varied systematically to determine how predictive performance depended on the number of qubits, circuit depth, number of embedded features, feature-embedding strategy, and amount of data used during QNN optimization.

Each experiment was repeated three times using different random seeds. These repetitions changed the parameter initialization of the single-qubit circuit to avoid a single initialization getting stuck in a local minium, while the parameters introduced upon adding subsequent qubits were initialized to zero as described above. In the data-fraction experiments, the seed also determined the subset of samples used to train the QNN. The reported metrics for each configuration correspond to the repetition with the best loss. Binary classification metrics were calculated using a probability threshold of 0.5, while AUC-ROC was used as the primary threshold-independent performance measure.

### Dependence on Circuit Architecture

We first examined the dependence of model performance on the number of qubits and data-reuploading layers. Increasing the number of qubits initially improved held-out predictive performance, but the improvement diminished as the circuit approached eight qubits (Figure 2A). Training performance continued to increase more readily than held-out performance, indicating that additional qubits primarily increased model capacity after the dominant generalizable structure of the data had been captured.

**Figure 1.**
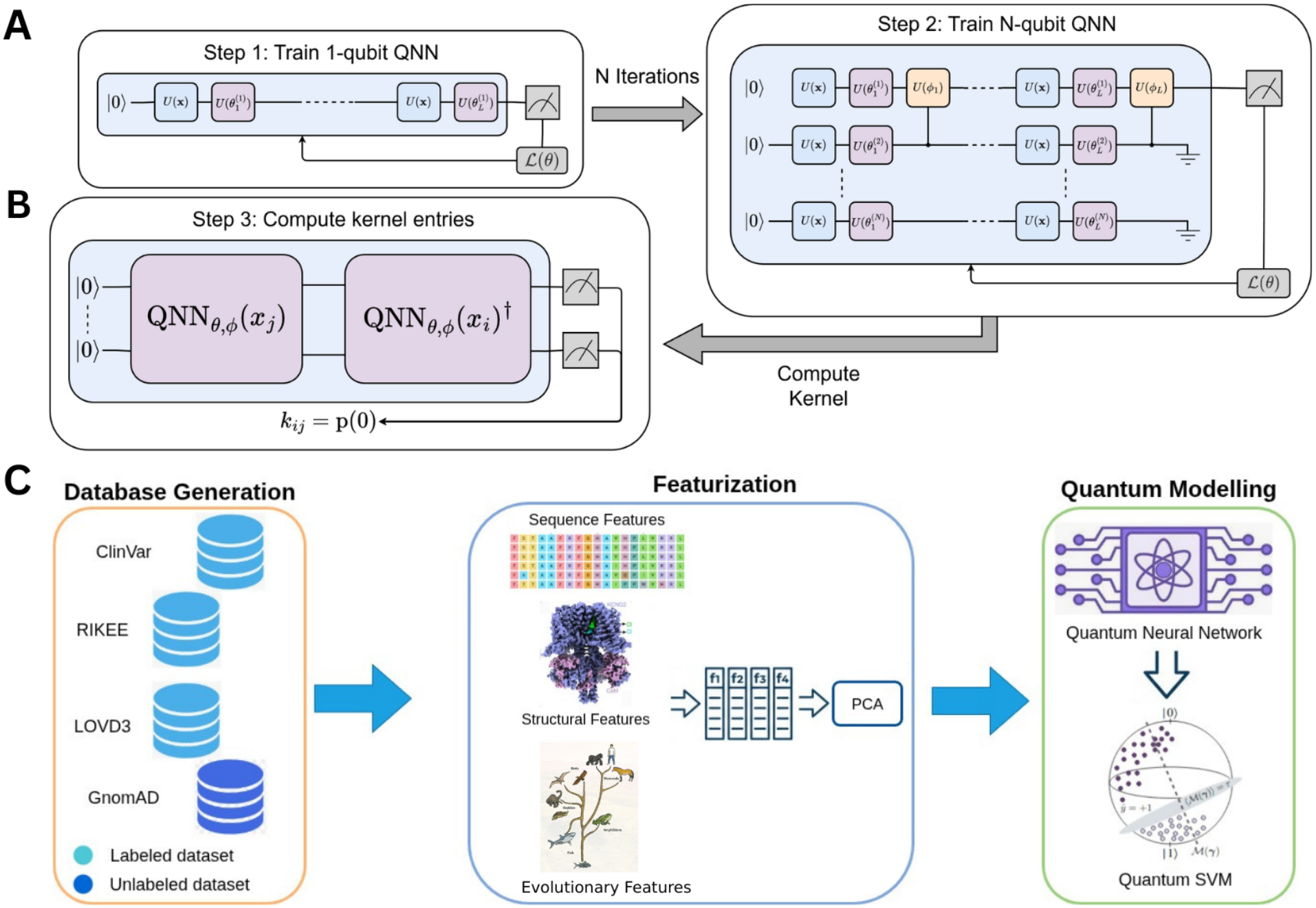
(A) Schematic representation of the QNN training procedure. The QNN circuit contains L layers of pairs of parameterized unitary transformations, the first parameterized by the data features, and the second by trainable parameters. The QNN is trained by iteratively adding qubits to the circuit, starting with a single qubit circuit which is trained to convergence, and sequentially adding additional qubits. After the first qubit, entanglement gates are added between adjacent qubits after each layer. (B) The EQK is trained by stacking the trained QNN circuit for a given datapoint i and its adjoint for another datapoint j, then applying the circuit to the zero state. The inner product between the resulting quantum state and the zero state provides the kernel entry *k*_*ij*_. The full kernel can then be used to train a support vector machine on the full training dataset. (C) Schematic representation of the workflow implemented in this work. Variant is collected from a variety of labeled clinical datasets (ClinVar, LOVD3, RIKEE) and an unlabelled dataset (gnomAD), which is labelled using heuristic criterions. Each variant is featurized using available structural and sequence based information about the mutation, and PCA is used to compress the size of the featurized representation of each variant. The top N principal components of this compressed representation are embedded into a quantum neural network using angle embedding. This QNN is trained to convergence and the trained circuit is then used to construct the quantum kernel of a support vector machine.

**Figure 2.**
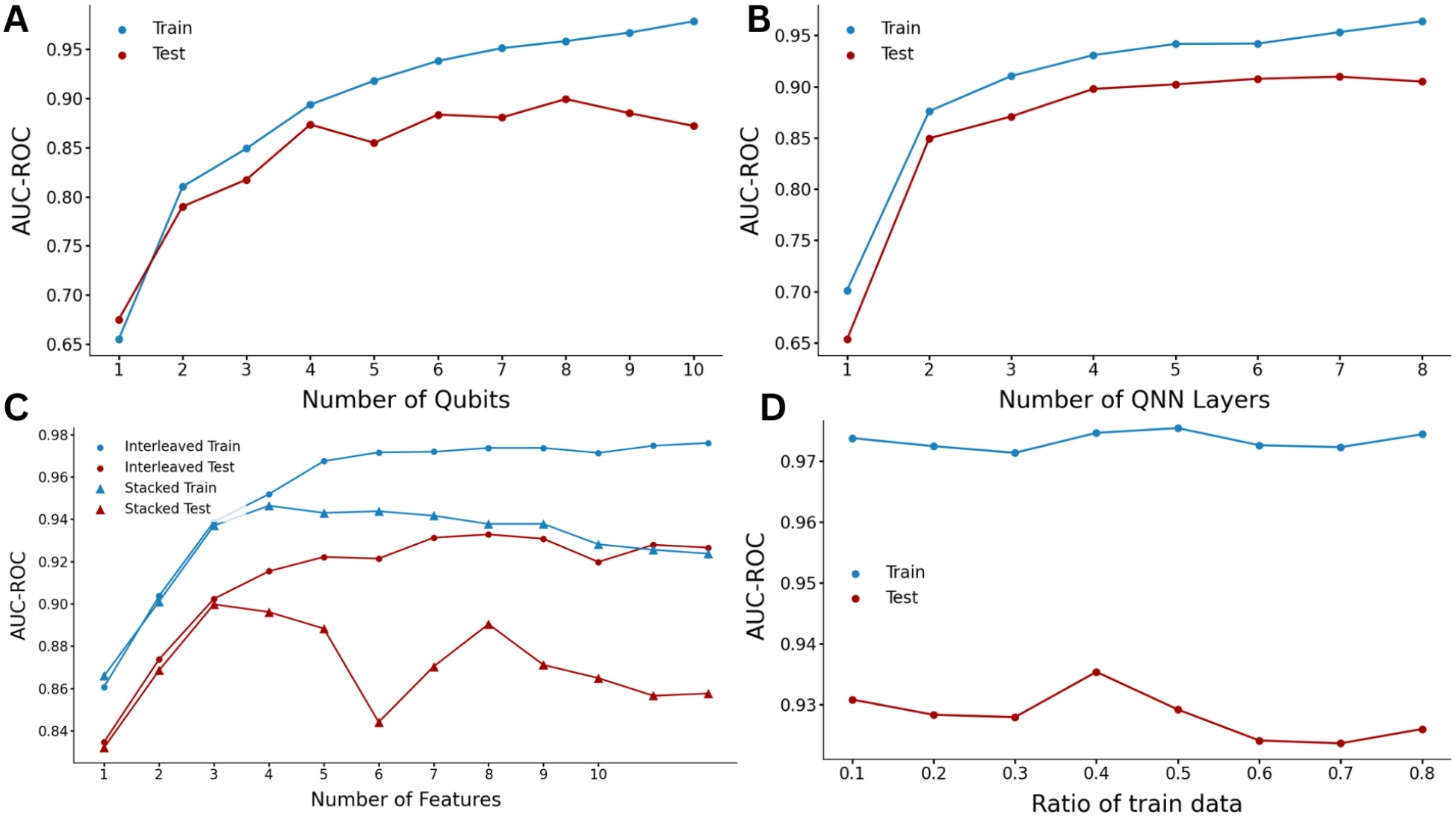
(A) Scaling of the quantum machine learning model with the number of qubits used in the QNN, for both the training and test datasets. (B) Scaling of the quantum machine learning model with the number of layers in the QNN, for both the training and test datasets. (C) Performance of different embedding strategies for embedding more than 3 features in the QNN, and their scaling as a function of the total number of embedded features. (D) Scaling of the quantum machine learning model with the fraction of the full training dataset used to train the QNN portion of the model, for both the training and test datasets. The full training data was used for the EQK portion of the model.

A similar trend was observed with circuit depth (Figure 2B). Increasing the number of data-reuploading layers improved performance for shallower circuits, whereas comparatively little additional improvement was observed beyond approximately six layers. These results indicate that the classification task benefits from increased circuit expressivity up to a moderate circuit size, after which additional depth or width produces diminishing returns.

These saturation trends should not be interpreted as evidence that larger circuits are generally unnecessary for variant-effect prediction. Rather, they indicate that increasing circuit capacity beyond this range provided limited additional benefit for the present KCNQ gene focused learning task.

### Number and Distribution of Embedded Features

We next investigated how performance depended on the number of PCA components supplied to the quantum circuit and on the strategy used to distribute those components across the circuit layers (Figure 2C). Two approaches were compared. In the stacked strategy, all retained components were encoded sequentially within every layer. In the interleaved strategy, consecutive component triplets were assigned to successive layers and cyclically re-uploaded across the circuit depth.

The interleaved strategy consistently produced higher held-out AUC-ROC values than the stacked strategy as the number of retained components increased. Performance improved as additional principal components were incorporated, but the gain diminished after approximately eight components. This behavior is consistent with the lower-ranked components containing progressively less useful signal, and redundant information becoming more difficult to encode effectively when many rotations are applied within the same layer.

The superior performance of interleaving suggests that distributing features across the circuit depth preserves the information contained in additional components more effectively than repeatedly stacking all corresponding rotations within each layer. The interleaved strategy was therefore used in the final model.

### Data Requirements for QNN Representation Learning

The most computationally demanding optimization stage is the training of the parameterized QNN. We therefore investigated whether the QNN required access to the complete training dataset in order to learn a useful quantum feature representation. For this experiment, the QNN was optimized using different fractions of the available training partition. Samples were selected randomly and stratified by class to preserve the benign-to-pathogenic ratio of the complete training data. Three repetitions were performed using different random seeds and, consequently, different parameter initializations and sampled subsets.

After QNN optimization, the resulting circuit parameters were fixed and the trained circuit was used to construct the quantum kernel over the complete training partition. Thus, only the data used during QNN optimization were reduced; the support vector machine was still trained using the full quantum kernel and all available training samples.

Held-out performance was comparatively stable even when the QNN was trained using only a fraction of the available data (Figure 2D). This indicates that the variational circuit can learn a sufficiently informative feature map from a representative subset, after which the complete dataset can be incorporated during kernel construction and SVM training. The result is relevant to larger-data applications because it suggests that the more difficult variational optimization stage need not necessarily scale with the complete number of available samples.

When attempting to train the full model using the same data subset, however, no such train-set size independence was observed, and performance reliably increased with the fraction of training data, suggesting that the amount of available data is still the fundamental bottleneck in this learning problem. Thus, this reduction applies specifically to QNN optimization and does not eliminate the computational cost of evaluating the full kernel matrix, which still grows with the number of samples used by the SVM. Nevertheless, separating the amount of data required for representation learning from the amount used for kernel-based classification may substantially reduce the cost of training the parameterized circuit.

Taken together, the architectural analyses show that performance becomes relatively insensitive to further increases in circuit width, circuit depth, and embedded feature count beyond moderate values. In contrast, the learned representation remains useful when QNN optimization is performed on a reduced but class-balanced subset. These observations suggest that an appropriately selected circuit can capture the dominant structure of the input representation without requiring either excessive circuit complexity or exposure of the QNN optimizer to the complete training dataset.

### Comparison with Classical Models

We finally compared the predictive performance of the quantum machine learning model against a set of standard classical machine learning approaches.

Across all evaluations (figure 3), the quantum model achieves performance comparable to that of the best-performing classical methods. In particular, it consistently reaches AUC-ROC values around 0.94, closely matching optimized classical approaches such as ensemble-based models. Although the quantum model did not show a consistent qualitative advantage over the classical baselines, the quantum model reliably ranks among the top-performing methods, demonstrating that its feature representation is sufficiently expressive to capture the relevant structure of the data. This demonstrates the feasibility of applying quantum machine learning methods in this domain and suggests that quantum feature mappings can encode meaningful representations of complex biological data.

**Figure 3.**
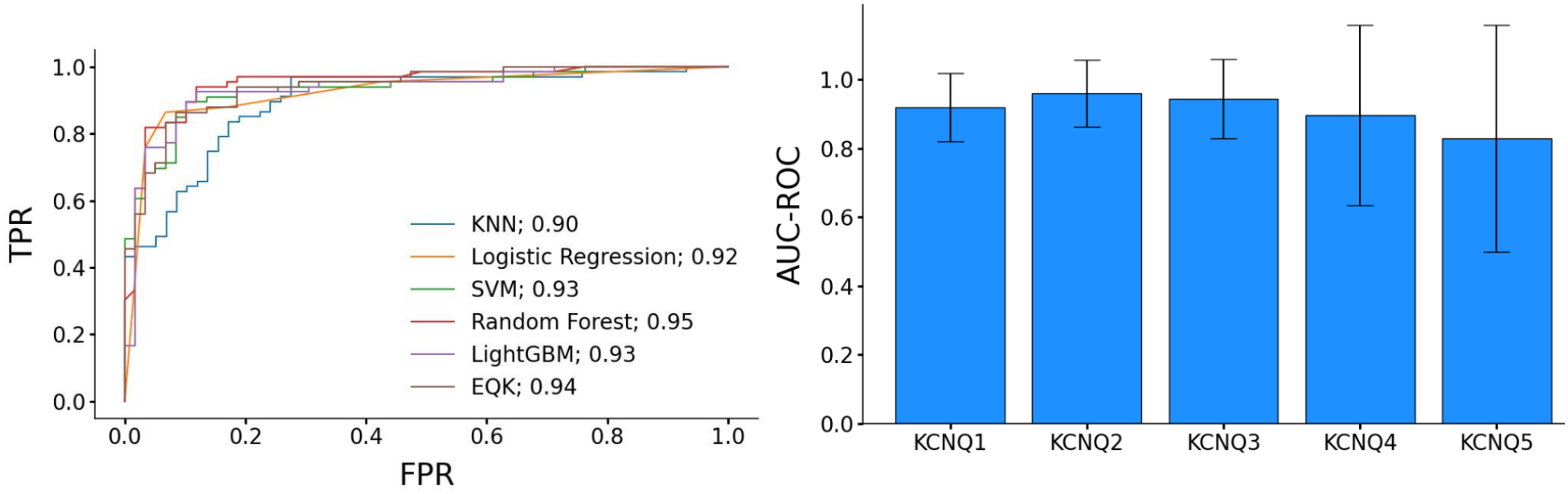
(A) ROC curves for the reference classical benchmarks and the quantum EQK model presented in this work. (B) AUC-ROC performance metrics for the quantum EQK model stratified into each of the genes in the KCNQ family.

Overall, these results indicate that predictive performance in this setting is largely governed by data limitations rather than model expressivity. While no clear advantage of the quantum model is observed in aggregate performance metrics, its ability to achieve competitive accuracy supports its use as a viable alternative framework. Future advances in dataset scale and quality, as well as improvements in quantum hardware, may help clarify the conditions under which quantum approaches can offer practical benefits for biological prediction tasks.

## Discussion

Predicting the pathogenicity of genetic variants remains a central challenge in computational genomics, particularly for genes such as the KCNQ family, where missense mutations are associated with a range of clinically significant phenotypes. Despite the growing availability of variant annotations, the number of confidently labeled examples remains limited, and the underlying relationship between sequence variation and functional impact is complex and multifactorial. These characteristics make pathogenicity prediction a representative example of a data-constrained learning problem in biology.

In this work, we explored the application of a quantum machine learning framework to this problem. The results demonstrate that this approach achieves predictive performance comparable to that of well-established classical machine learning models across multiple evaluation metrics. In particular, the quantum model matches the performance of optimized classical baselines such as random forests and gradient boosting models, indicating that the learned quantum representation is sufficiently expressive to capture the relevant structure of the data.

A key observation from our analysis is that model performance exhibits consistent saturation behavior across a range of architectural parameters, including circuit depth, number of qubits, and number of embedded features, while increasing the size of the training dataset does lead to improved performance, which supports the interpretation that the bottleneck for modeling this family of genes lies in the data rather than in the modeling approach. Our exhaustive hyperparameter exploration also showcases that the performance of these quantum model is more sensitive to hyperparameter choices compared to classical models, but this analysis provides some perspective on which choices are adequate for the kind of learning problem tackled in this work.

This study shows that quantum machine learning can be successfully applied to the prediction of variant pathogenicity, achieving performance on par with established classical methods in a challenging gene family. While no clear advantage of the quantum approach is observed at present, the results provide evidence that quantum feature representations are viable for this class of problems. As both datasets and quantum technologies continue to evolve, further work will be needed to determine whether and when quantum methods can offer meaningful improvements in biological prediction tasks

## Data and Software Availability

The input files, structure files and other supporting material necessary to reproduce the results reported in this study have been deposited in a public repository at GitHub:

## Acknowledgements

Part of this material is based upon work supported by the National Science Foundation under Grant No. DBI-1548297 (S. C.).

